# The Effect of Radiotherapy on Head and Neck Cancer Cell Lines Following Exposure to Heat

**DOI:** 10.1101/2020.11.24.385070

**Authors:** Ben Atkinson, Jamie Wilson

## Abstract

Fanconi anaemia (FA) is a rare, recessive, genetic disorder characterised by a predisposition to cancer. Patients with FA are 700 times more likely to develop head and neck squamous-cell carcinomas including oral epithelial dysplasia (OED), a potentially malignant disorder of the oral mucosa. This increased likelihood suggests that the molecular mechanism responsible for dysplastic transformation may involve defects in the FA pathway. In this study, the significance of ataxia telangiectasia and Rad3-related protein (ATR), which is responsible for homologous recombination repair (HRR), was investigated. ATR protects against mutations in both normal squamous cells and cancer cells by inducing HRR, and thus is associated with radiotherapy resistance. This investigation was designed to study the effects of heat on DNA repair specific to the FA pathway. Western blotting was carried out to determine whether heat affects the ATR pathway, followed by survival assays to determine the viability of cells after heat treatment and then compared to cells treated with a specific ATR inhibitor. This project aims to discover whether heat could be used as a non-invasive treatment to increase the sensitivity of tumour cells towards radiotherapy leading to an improved treatment plan for patients suffering from head and neck cancers.

## Introduction

Maintaining the integrity of DNA in an unchanged and intact state is crucial for many biological functions from replication to the transfer of genetic information from generation to generation. However, DNA is highly susceptible to damage. Mutations have the potential to cause alterations to protein structure or even result in a total loss of expression [1]. The body has developed multiple mechanisms to detect, repair and even destroy damaged DNA, often resulting in apoptosis if the damage is too severe. The Fanconi Anaemia pathway is one such repair pathway. Patients with Fanconi anaemia disease are disproportionately affected by head and neck cancer suggesting this pathway plays a crucial protective role for cell survival. We specifically looked at how the phosphorylation of ataxia telangiectasia and Rad3-related protein (ATR) in the FA pathway initiates homologous recombination repair (HRR). It has previously been evidenced that patients suffering from Fanconi anaemia have a mutation in proteins such as ATR, hindering HRR, which accumulates mutational errors by repairing through the less precise non-homologous end joining (NHEJ) techniques and thus giving rise to epithelial dysplasia and cancers.

Radiotherapy is used alongside chemotherapy in the treatment of cancers. Cancer cells exposed to radiation ineffectively repair irradiation-induced DNA damage compared to normal tissue cells [2]. Studies have shown that increasing the dosage of radiation does not increase its efficacy. In fact, the occurrence of side effects, such as ulceration, mucositis, sialadenitis found in head and neck cancer patients is proportional to the dosage. Radiation initiates severe DNA damage in the form of a double strand break (DBS) which if present in either normal cells or cancer cells can cause the cell to become inviable if not repaired. Through a protein cascade, ATR kinase is activated which initiates HRR for effective cell survival. A breakdown in this pathway induces the rate of NHEJ which has the potential to lead to mutations and the development of carcinomas. ATR remains active in cancer cells to repair DSBs and if these are not repaired the cell is forced to undergo apoptosis. Inhibiting ATR in the FA pathway has the potential to evolve current cancer treatments by preventing the repair of DSBs.

It has been hypothesised that elevating the cellular temperature can aid in tumour regression by direct inhibition of ATM, an important DNA repair enzyme [3]. Due to the close relationship of ATM and ATR it stands to reason both would be affected by mild hyperthermic stress. We designed this experiment to show the effect heat has on the expression of ATR in response to DNA damage and how this affects the survival of various squamous cell carcinoma cell lines. It is our hope that we are able to show that it is possible to use heat as a non-invasive treatment to increase the efficacy of radiotherapy and subsequently leading to an improved treatment plan for patients undergoing radiotherapy for head and neck cancers.

## Materials and methods

### Cell culture

Three cancer cell lines, UMSCC6, UMSCC74A and UMSCC47, were used throughout this study. They were routinely cultured in Dulbeccos’s Modified Eagles media (Sigma, Poole Dorset) supplemented with 10% foetal calf serum and 1% penicillin-streptomycin, and maintained at 37 °C, 5% CO_2_. Cell lines were transfected with a hygromycin resistance cassette and a neomycin flanking cassette as described previously.

To mimic radiotherapy, cells were irradiated at 2 Gy for 60 seconds. To induced mild temperature hyperthermia cells were warmed to 39°C in an incubator for 15 minutes. ATR inhibition was carried out using (ATR inhibitor M6620 (VX-970)) at 100 nM for 24hrs.

### Survival assay

Colony formation assays were using to calculate cellular viability following treatment with each of the individual or combination treatments. Briefly, 1000 cells were seeded into a 10 cm dish and treated with their respective treatments. After two weeks, cells were fixed in 10% formalin and stained with crystal violet for 5 minutes at room temperature. The number of colonies were then quantified by eye.

### Western Blotting

Levels of ATR and ATR S428 were quantified by Western Blotting. Protein samples were collected from UMSCC47 cells following radiation, hyperthermia (39°C), ATR inhibitor (100nM), or both combined. Each sample was loaded onto a 15% SDS acrylamide gel and ran over night at a constant 60V. Proteins were transferred to nitrocellulose membrane by at 350 mA and once complete, blocked using LiCor blocking buffer at room temperature for 1 hour. Specific antibodies against ATR 1:1000 (Atlas antibodies Cat no. HPA054320) and ATR S428 1:2000 (cell signalling Cat no. 2853s) were applied and incubated for 2 hours at room temperature. Following washing with TBS-tween, secondary antibody (Invitrogen Cat no. A32735) was added, and membranes incubated in the dark, at room temperature for 2 hours. Membranes were then developed using a fluorescent scanner.

### Statistical tests

All data has been analysed using a two-way unpaired t-test. For the purpose of this study, a P value of <0.05 is deemed significant.

## Results

### Western Blotting

In order to determine whether mild hyperthermia can disrupt the ATR pathway and sensitise patients to radiotherapy, cells were irradiated following exposure to mild hyperthermia or an ATR inhibitor and protein expression for total ATR and phospho-ATR was quantified by a western blot. Phosphorylation of S428 was examined as this phosphor-site has been shown to be required for the activation of the downstream ATR kinase pathway. We have shown that expression of total ATR remains constant independent of treatment group. However, we have demonstrated that following mild temperature hyperthermia, pharmacological ATR inhibition or a combination of the two, there is a decrease in ATR S428, suggesting that mild hyperthermia may prevent activation of the ATR pathway.

**Figure 1.**
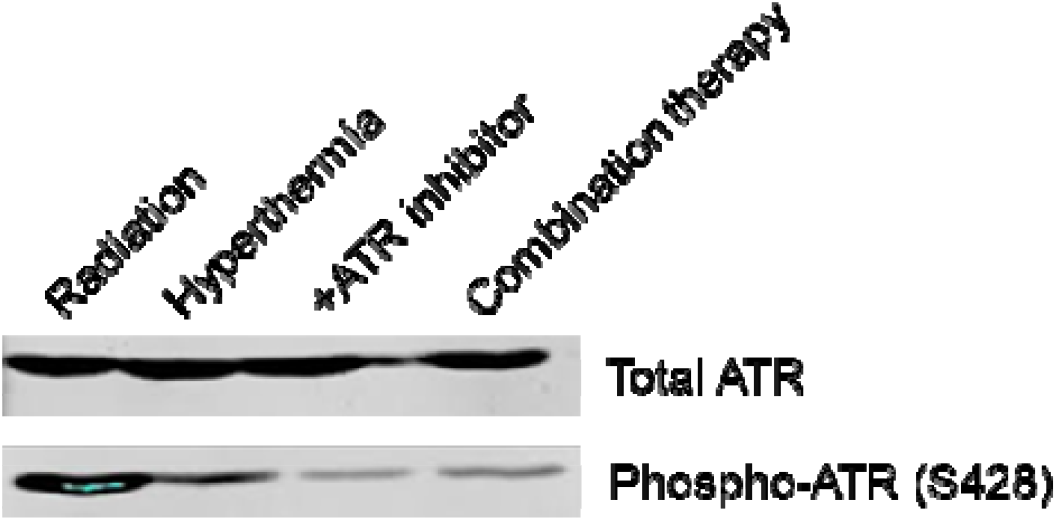
Immuno blot using specific antibodies for the ATR and phosphorylation site S428 ATR on proteins extracted from UMSCC47 and treated as follows, lane 1 radiation only, lane 2 heat (39°C), lane 3 inhibition (100nM), lane 4 heat and inhibition. A sing ATR antibody showing ATR expression level after all treatments. B using ATR S428 specific antibody showing a decrease in expression after treatment.

### Clonogenic Survival Assay

Having established that heat reduces phosphorylation of ATR, we next sought to further investigate whether mild hyperthermia could affect cellular survival *in vitro*. To do this, clonogenic survival assays were carried out in four head and neck cell lines following treatment with radiation, mild hyperthermia, ATR inhibition or a combination therapy.

For this work the control group was defined as cells treated with radiation only to induce DSB’s, stimulating ATR associated repair mechanisms. Treatment with heat and ATR specific inhibitors shows a reduction in survival, with heat only cells having between 78-83% survival (Fig 2). Analyses showed there to be no significant difference in the survival rates of cells treated with heat compared to those treated with inhibitor (P>0.05) (Fig 2). Likewise, no significant difference was found between UMSCC6/UMSCC47 cells exposed to heat and cells exposed to heat and inhibitor (P>0.05) along with and cells exposed to a combination of heat and inhibitor(P>0.05) (Fig 2). UMSCC74A survival was shown to be the same for all groups apart from cells that underwent heat treatment when compared to Inhibitor. This comparison did in fact show a significant difference (P<0.05).

**Figure 2.**
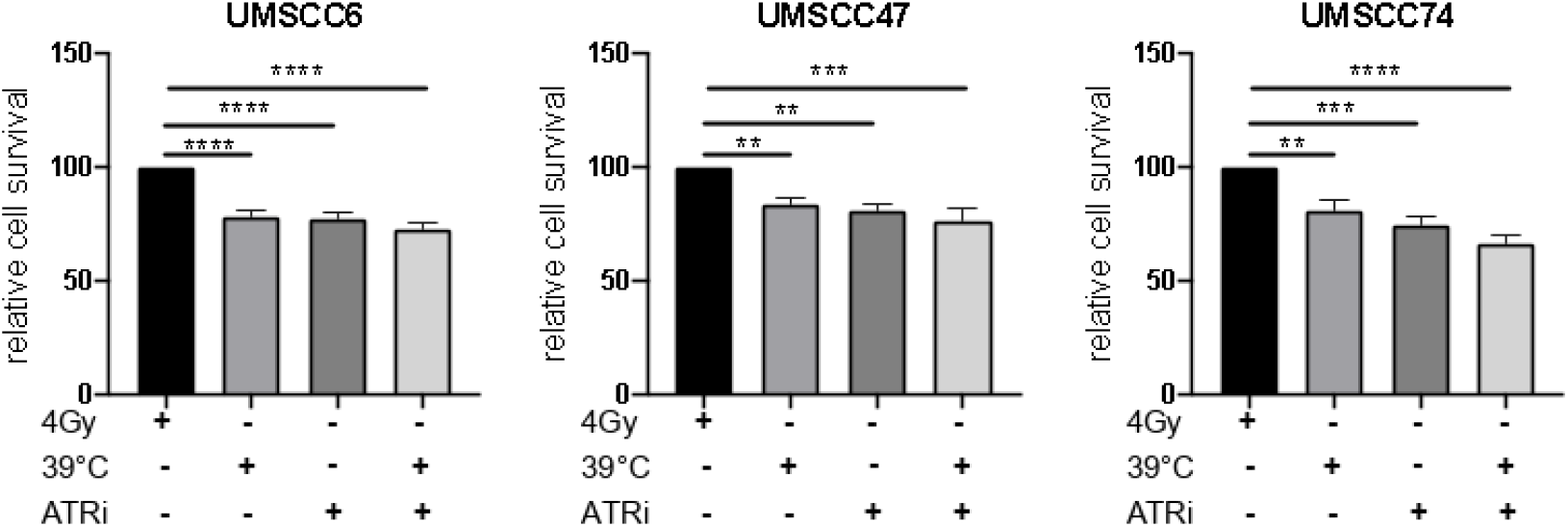
Results of clonogenic survival assays of 3 cell lines (UMSCC6, UMSCC47, UMSCC74A) after treatment with radiation only heat (39°C), ATR inhibitor (100nM), and both combined.

## Discussion

The aim of this study was to determine the effect of mild hyperthermia on HRR, in the context of ATR kinase. We have demonstrated that although total ATR protein expression is not affected by hyperthermia, phosphor-ATR (S428) is decreased as a result of heating cells to 39 °C. The decrease in phosphorylation, although not as large as that induced by pharmacological intervention, is notable. ATR S428 is a key protein involved in the repair of DSBs by activating HRR. However, in cells treated with heat, ATR inhibitor and both combined before being exposed to radiation showed a reduction in the level of ATR S428 present in the cell. The inhibitor is preventing the phosphorylation of ATR, thus decreasing the amount of ATR S428 present to activate HRR. The lack of ATR S428 seen after heat and both combined suggests heat is having the same effect on ATR as the inhibitor. This supports the theory that hyperthermia can inhibit the activation of ATR. The level of ATR S428 following treatment with heat was consistent with the level seen in cells after treatment with both heat and inhibitor suggesting the effect of heat and inhibitor is not additive.

To investigate a functional effect of decreased ATR phosphorylation, we studied clonogenic survival following cell treatment. Our data demonstrates that, when compared to radiation controls, cells exposed to a hyperthermia-ATR inhibitor combination therapy had a statistically significant reduction in cellular survival. Our data suggests that the combination therapy has an additive effect at the functional level and is superior to inhibitor and hyperthermia monotherapies. When combined with data from figure 1, this data suggests that the combination therapy may be affecting other pathways outside of the ATR signalling cascade.

The clinical aim of this study was to evaluate the efficacy of a radiotherapy-hyperthermia combination therapy to inhibit the mechanisms involved in DSB repair. The significant reduction in survival of cells treated with heat prior to being exposed to radiation strongly suggests that heat can sensitise cancer cells to the effects of radiation.

Throughout this study an ATR inhibitor at a concentration of 100nM was used. As seen from the western blots there was still some residual ATR S428 in cells treated with the inhibitor, suggesting there was not a total inhibition taking place. This allowed us to compare the effect of heat more evenly to cells treated with inhibitor, as a similar level of inhibition was seen in cells treated with heat.

## Abbreviations

ATM: Ataxia-telangiectasia mutated
ATR: Ataxia-telangiectasia and Rad3 related
HRR: Homologous recombination repair
DNA: Deoxyribonucleic acid
NHEJ: Non-homologous end joining
DSB: Double strand break

## Acknowledgements

Thank you to the North West Cancer research centre and the University of Liverpool for their support throughout my time there.

